# Spotting the path that leads nowhere: Modulation of human theta and alpha oscillations induced by trajectory changes during navigation

**DOI:** 10.1101/301697

**Authors:** Amir-Homayoun Javadi, Eva Zita Patai, Aaron Margois, Heng-Ru M. Tan, Darshan Kumaran, Marko Nardini, Will Penny, Emrah Duzel, Peter Dayan, Hugo J. Spiers

**Affiliations:** Institute of Behavioural Neuroscience, University College London, London, UK; School of Psychology, University of Kent, Canterbury, UK; Institute of Cognitive Neuroscience, University College London, London, UK; Google Deepmind; Department of Psychology, Durham University, UK; School of Psychology, University of East Anglia, UK; Institute of Cognitive Neurology and Dementia Research (IkND), University Hospital Magdeburg, Germany; Gatsby Computational Neuroscience Unit, ucl, London, UK

## Abstract

The capacity to take efficient detours and exploit novel shortcuts during navigation is thought to be supported by a cognitive map of the environment. Despite advances in understanding the neural basis of the cognitive map, little is known about the neural dynamics associated with detours and shortcuts. Here, we recorded magnetoencephalography from humans as they navigated a virtual desert island riven by shifting lava flows. The task probed their ability to take efficient detours and shortcuts to remembered goals. We report modulation in event-related fields and theta power as participants identified real shortcuts and differentiated these from false shortcuts that led along suboptimal paths. Additionally, we found that a decrease in alpha power preceded ‘back-tracking’ where participants spontaneously turned back along a previous path. These findings help advance our understanding of the fine-grained temporal dynamics of human brain activity during navigation and support the development of models of brain networks that support navigation.

## Introduction

The neural circuits that support spatial navigation have been extensively studied in rodents and humans (Epstein, Patai, Julian, & Spiers, 2017b; Hartley, Lever, Burgess, & O‘Keefe, 2013). However, little is known about the neural processing that occurs when detours or shortcuts are taken to navigate to goals (Epstein, Patai, Julian, & Spiers, 2017a; Spiers & Gilbert, 2015). This is surprising, given that shortcuts and detours are key evidence of the use of a cognitive map of the environment to support behavior (O‘Keefe & Nadel, 1978; Tolman, 1948). A number of human neuroimaging studies using virtual reality have reported increased activity in prefrontal regions when a forced detour is required (Howard et al., 2014; Iaria, Fox, Chen, Petrides, & Barton, 2008; Maguire et al., 1998; Rauchs et al., 2008; Rosenbaum, Ziegler, Winocur, Grady, & Moscovitch, 2004; Simon & Daw, 2011; Spiers & Maguire, 2006; Viard, Doeller, Hartley, Bird, & Burgess, 2011; Xu, Evensmoen, Lehn, Pintzka, & Håberg, 2010). However, no previous neuroimaging study has examined the neural response evoked by shortcuts specifically and in isolation from detours (Ribas-Fernandes et al., 2011). In rodents, electrophysiological recording of hippocampal place cells has revealed ‘remapping’ in response to the changes in barriers that induced detours or shortcuts (Alvernhe, Save, & Poucet, 2011; Alvernhe, Van Cauter, Save, & Poucet, 2008; Poucet, Thinus-Blanc, & Chapuis, 1983). However, these studies examined neural coding of the new maze geometry (e.g. place cell remapping) rather than the event-related responses evoked by the changes to the maze. Moreover, despite extensive research on the neural oscillations that arise during navigation in rodents, few studies of human navigation have examined neural oscillations or evoked responses at a fine-grained time-scale in relation to the spatial processing involved (Cornwell, Johnson, Holroyd, Carver, & Grillon, 2008; Kaplan et al., 2014; Vass et al., 2016), thus limiting cross-species comparisons of the underlying neural mechanisms during navigation in dynamic environments.

In order to investigate neural responses and oscillations evoked by shortcuts and detours we recorded magnetoencephalography (MEG) while participants navigated to remembered goal locations in a virtual desert island containing paths constrained by lava flows (Figure 1, and Methods). One day before testing, participants learned the layout and location of 20 hidden objects in the environment. During MEG recording the lava could shift position, opening new paths (Shortcuts and False Shortcuts) or closing off paths (Detours), with one change type per trial (i.e per each object search period). While Shortcuts provided new paths that, if taken, would shorten the path distance to a goal, False Shortcuts would lead to longer paths to a goal (see Fig. 1B); they could either (False Shortcut Toward) appear to lead in the direction of the goal (a characteristic shared with Shortcuts) or appear to lead away from the goal (False Shortcut Away). As Shortcuts and False Shortcuts were visually identical, participants had to have an accurate cognitive map of the maze layout in order to choose whether to take the opening in the lava or not. To test whether the amount of change in the distance to the goal affected neural responses, lava changes associated with Shortcuts and Detours could lengthen or shorten the optimal remaining path by either 4 or 8 maze segments. Furthermore, subjects would sometimes spontaneously ‘Back-track’, by returning along their previous paths. Back-tracking could either be correct (bringing the subjects closer to the goal) or incorrect (taking them further away). We compared event-related fields and oscillatory markers that arose immediately after the three types of terrain change: Shortcuts, False Shortcuts and Detours, and prior to spontaneous Back-tracks.

**Figure 1:**
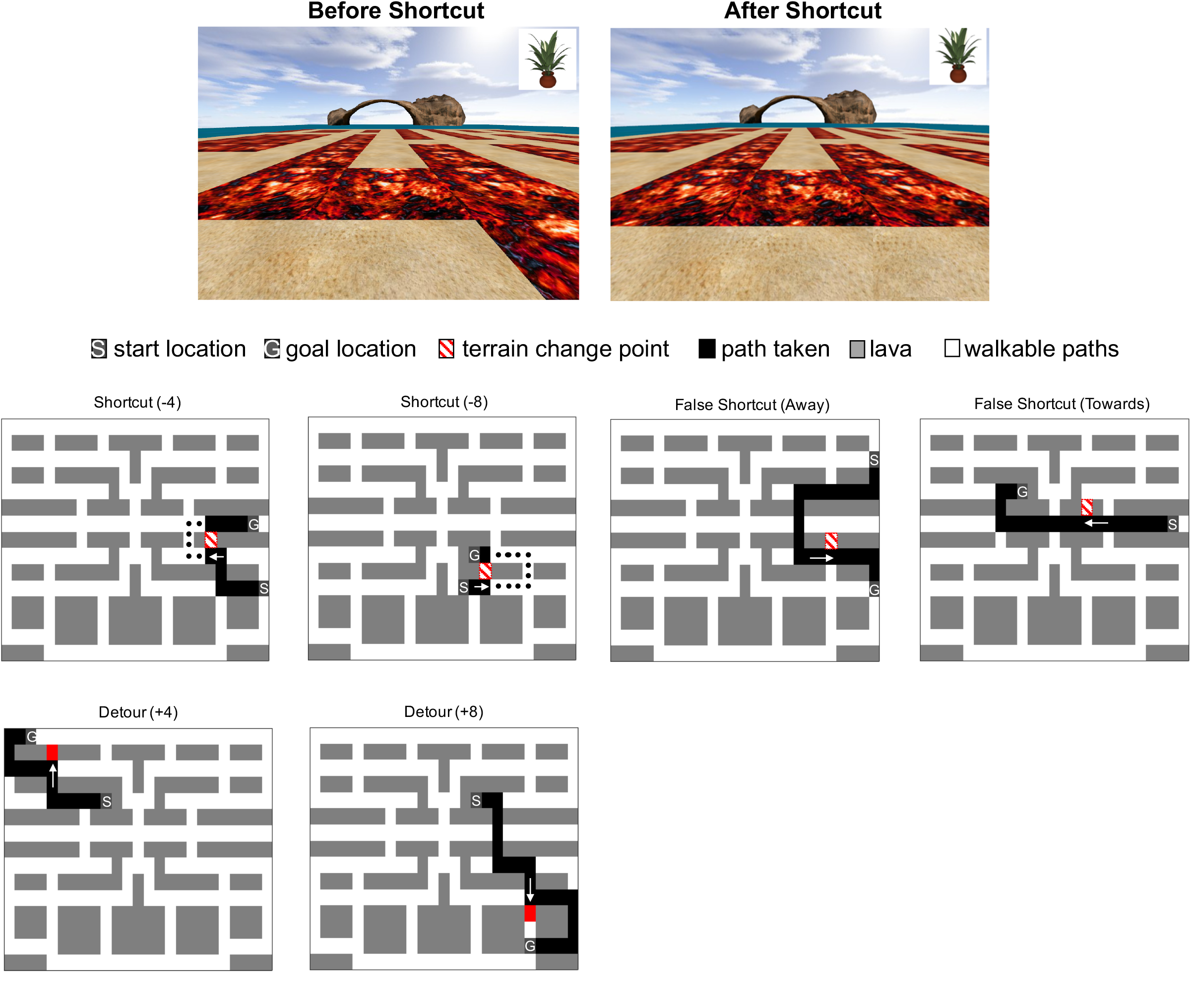
Lava World. Top: Example view of test environment before (left) and after (right) a change (in this case, inducing a shortcut). During each trial, subjects had to navigate along the sand using controls to move forward, left/right and backward to get to the current goal object (shown in the top right corner, counterbalanced left/right across participants) without stepping on the red ‘lava’. A distal cue is visible (arch), and 3 others were located at the other cardinal directions. The spatial location of the target was hidden. On each trial, one lava segment shifted to reveal new paths (Shortcuts or False Shortcuts), or could close an existing path (Detours). Bottom: Examples of change types.

## Results & Discussion

### Behaviour

We found that certain types of terrain change resulted more in incorrect choices at the change point, in other words, more suboptimal route choices (F(1,23)=13.04,p<0.001), a trend towards the magnitude of change in distance bein related to suboptimal choice (F(1,23)=3.61,p=0.07), and a significant interaction (F(1,23)=8.87,p=0.007). Long Detours (+8 maze units) resulted in more suboptimal choices compared to all other conditions except False Shortcuts Towards (all remaining t(1,20)<-3.02,p<0.006). Except for Long Detours (t(1,23)=0.24,p=0.8), False Shortcuts Towards resulted in more suboptimal choices than all other conditions (all t(1,23)<3.07,p<0.005). See Table 1, and Extended Data Table S1 for comprehensive t-tests. There was also a significantly higher propensity (i.e., lower criterion) to take False Shortcuts Towards the goal, compared with False Shortcuts Away from the goal (t(1,23)=-7.01, p<0.001).

For further scrutiny we examined the number of extra steps (maze units, see Fig. 1 Lower panel) as a proportion of the total (new) number of optimal steps after the terrain change events. We found Shortcuts resulted in the largest proportion of steps off-route (main effect of type: F(1,23)=35.3,p<0.001, see Table 1), despite overall more steps taken off-route in the Long Detour and Lures Towards condition Some of these extra steps were due to participants turning around, i.e., “Backtracking” (overall, 18% of the extra steps were such back-tracking events). These Back-tracking events were more common in the False Shortcuts Towards the goal and the Detours (+8) condition (both compared to all other events t(1,23)>2.1, p<0.046, but not different from each other (t(1,23)=1.0,p=0.32f see Extended Data Table S2 for details)f it was specifically the longer Detours that led to backtracking (interaction type × magnitude, F(1,23)=16.2,p<0.001). Moreover, we calculated the ratio of correct, compared to incorrect, back-tracking trials (“correct” is defined as a trials in which back-tracking would actually bring the participant closer to the goal), and found that overall 80% (±1% s.e.m, range: 50-100%) of back-tracking events were correct or optimal, and occurred equally frequently across all conditions (F(1,5)=0.27,p=0.9). Thus, in the majority of Back-tracking events participants became aware that they were heading in the wrong direction and spontaneously decided to turn around.

**Table 1:** Behavioural Results Summary [mean (±s.e.m)]

|  | **Detour**<br>**(+8)** | **Detour**<br>**(+4)** | **Shortcut**<br>**(-8)** | **Shortcut**<br>**(-4)** | **False**<br>**Shortcuts**<br>**Towards** | **False**<br>**Shortcuts**<br>**Away** |
| --- | --- | --- | --- | --- | --- | --- |
| **Correct Route**<br>**Choice [%]** | 70.2($\pm 4.5$) | 82.2($\pm 2.3$) | 84.8($\pm 3.4$) | 84.2($\pm 2.8$) | 71.1($\pm 4.2$) | 87.1($\pm 3.1$) |
| **Extra Steps**<br>**[proportion]** | 1.2($\pm 0.5$) | 1.3($\pm 0.5$) | 2.0($\pm 1.7$) | 1.9($\pm 1.0$) | 1.4($\pm 0.8$) | 1.6($\pm 1.4$) |
| **Back-tracking [#]** | 3.5($\pm 0.6$) | 2.1($\pm 0.4$) | 1.1($\pm 0.3$) | 1.9($\pm 0.4$) | 3.0($\pm 0.5$) | 1.8($\pm 0.5$) |
| **Correct**<br>**Backtracking [%]** | 81.3($\pm 4.5$) | 82.5($\pm 4.0$) | 81.3($\pm 7.3$) | 66.7($\pm 21.1$) | 79.5($\pm 3.9$) | 72.2($\pm 18.1$) |

### Electrophysiological Indices of Navigation

#### Shortcuts vs False Shortcuts

While visually identical, the comparison of Shortcuts and False Shortcuts (Towards and Away) allow us to investigate how the brain responds to changes in the environment that result in different values of outcome, i.e. real shortcuts are useful. False Shortcuts should be processed differently from Shortcuts if participants have an accurate memory of the layout of the maze and are able to differentiate between useful and not useful openings in the lava. When examining the wait period after a change point (a four second delay after the change in lava during which participants had to decide which route to take), we found that Shortcuts had significantly different event-related fields from both types of False Shortcut (Shortcuts vs False Towards: positive cluster p=0.04, time: 180-540ms, sensor distribution: central-posterior, and negative cluster p=0.014, time: 660-1000ms, sensor distribution: bilateral temporal-frontal, Figure 2A; Shortcuts vs False Away: p=0.008, time: 460-880ms, sensor distribution: right frontal-temporal, Figure 2B). Moreover, False Shortcuts Towards the goal were different from False Shortcuts Away from the goal (p=0.008, time: 460-880ms, sensor distribution: right frontal-temporal, Figure 2C). To confirm that these effects were not driven by changes in eye-movement activity, we conducted paired t-tests and found that saccadic behaviour (as measured by variance in the saccade components derived from the ICA, see Methods) was not significantly different between these conditions (Shortcuts vs False Shortcut Toward, t(1,23)<1.5,p>0.1, Shortcuts vs False Shortcut Away, t(1,23)=1.5,p=0.07), False Shortcuts Towards vs Away t(1,23)=0.26, p>0.1). Thus, rapidly after a terrain changes (as early as 180ms) neural processing distinguishes between potential useful new paths from those that will be detrimental in reaching the goal, and after around half a second distinguish two different types of false shortcut.

Prior research has indicated that oscillations at theta frequencies are involved in navigation and spatial memory (Bohbot, Copara, Gotman, & Ekstrom, 2017; Buzsáki, 2005; Chakravarthy & Balasubramani, 2013; Cornwell et al., 2008; Eschmann, Bader, & Mecklinger, 2018; Hartley et al., 2013; Hasselmo, Hay, Ilyn, & Gorchetchnikov, 2002; Hasselmo, Hinman, Dannenberg, & Stern, 2017; Jocham et al., 2014; Kaplan et al., 2014, 2012; Mohan et al., 2016; Snider, Plank, Lynch, Halgren, & Poizner, 2013). We therefore examined this frequency band (3-7Hz). Shortcuts led to significantly increased theta power compared to both types of False Shortcut for nearly the whole duration of the epoch (4s) after the change point (Shortcut vs False Shortcut Towards: 50-2140ms with a frontal-central distribution; Shortcut vs False Away: 0-3160ms with a bilateral frontal temporal distribution, see Figure 3A). We also found increased theta power over right parietal-temporal sensors for False Shortcuts Towards the goal compared to Away from the goal, starting late after the change point over left parietal sensors (2000-3000ms, p=0.03) (Figure 3A). There was no strong relationship between behavioural accuracy and the theta response (Shortcut vs False Shortcut Towards r=.34,p=0.099; Shortcut vs False Shortcut Away r=.19,p=0.37; False Shortcuts Towards vs Away r=.36, p=0.085;), indicating the changes in theta response are not a simple function of behavioural choice or difficulty.

**Figure 2:**
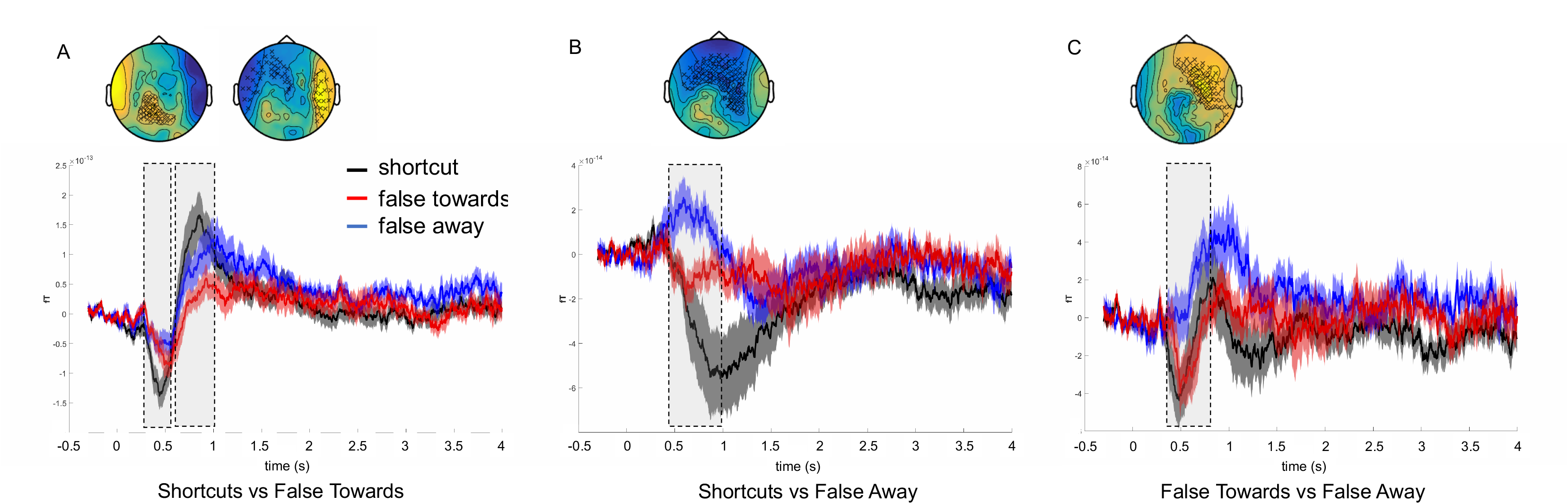
Event-related field changes to Shortcuts and False Shortcuts. We found significant differences in the ERFs between **A)** Shortcuts vs False Towards, positive cluster 180-540ms, and negative cluster p=0.014, time: 660-1000ms **B)** Shortcuts vs False Away 460-880ms and **C)** False Shortcuts Towards vs Away from the goal between 460 – 880ms, after the onset of the change point (opening in the lava). Displayed in each panel is the topography of the difference between the conditions with the significant sensors marked by x’s. The plotted ERF is the average (±s.e.m.) over the significant sensors, with the significant time-period highlighted with dashed boxes. All results are corrected for multiple comparisons using cluster-based permutation tests. (Plots of the responses in Detours are presented in Fig S1).

#### Detours vs Shortcuts and Effects of Change Magnitude

Due to eye-movements, we cannot explicitly rule out the effects between conditions are not contaminated by differences in saccadic variance. Please see Supplemental Results for details.

**Figure 3:**
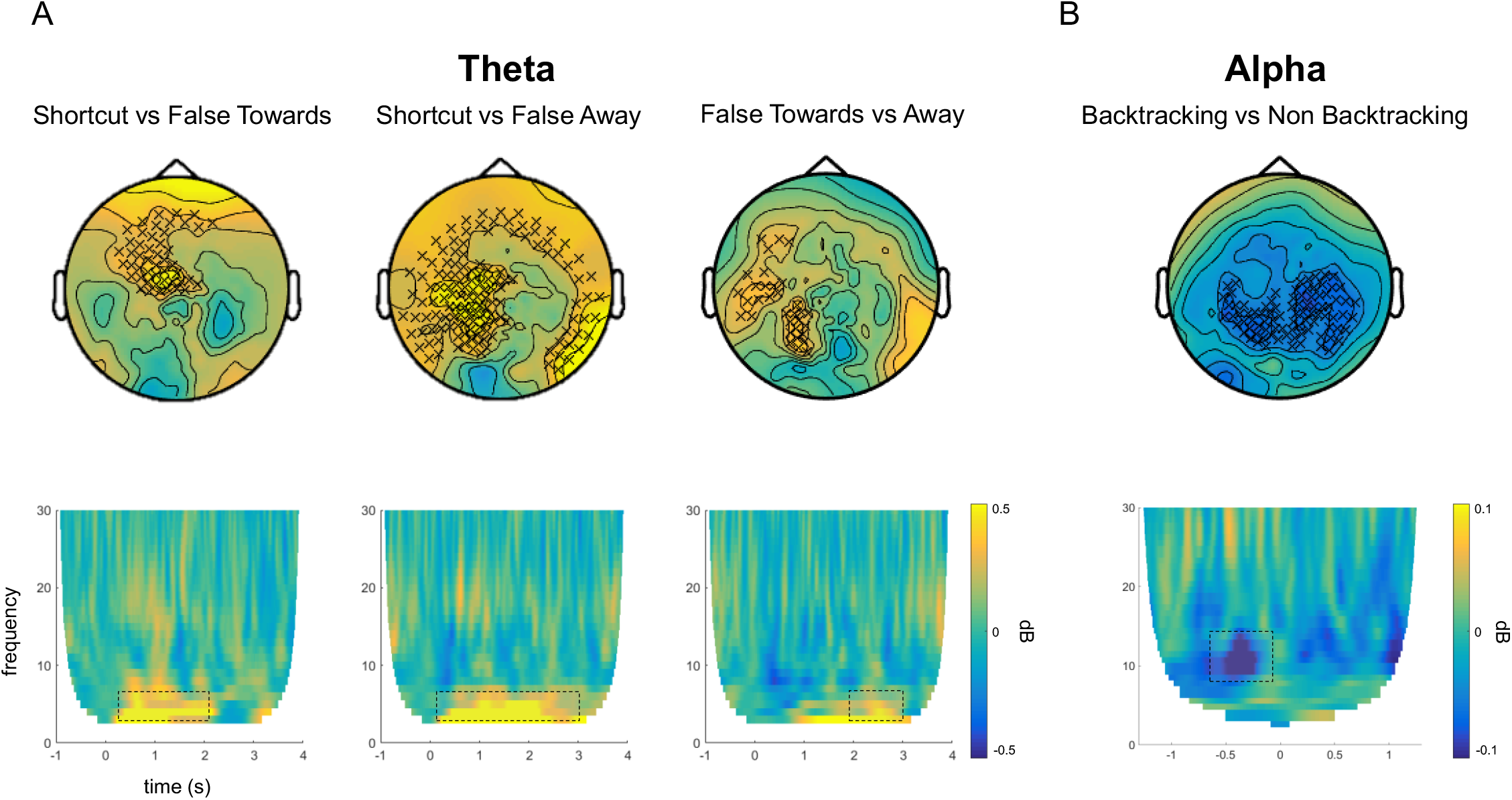
Distinct time-frequency markers for processing False Shortcuts and spontaneous Back-tracking. **A)** We found increased activity in the theta band when comparing Shortcuts to both types of False Shortcut (Towards: 50-2140ms; Away: 0-3160ms) as well as between False Shortcuts Towards and Away from the goal (2000-3000ms). **B)** Preceding spontaneous path changes, i.e. Backtracking, we found decreased alpha power over posterior sensors (−700 to −150ms before the Backtracking event). This effect was specific to the alpha band (no significant effects for theta or beta band in the same period). Top panels show topographies of significant sensors. Bottom displays time-frequency plots of the data A) combined over the three significant sensors that overlapped in all three condition comparisons and B) from all significant sensors, showing the specificity of frequency bands as well as temporal profile of condition differences (significant frequency/time-period highlighted with dashed boxes).

#### Backtracking vs Non-Backtracking

Next we investigated time points in which participants spontaneously decided to turn around (‘Back-tracking’). We compared the neural signatures of these events to equivalent periods in which participants did not backtrack (defined as steps that were at the same relative point in steps along another route, see Methods for details). We considered that Back)tracking would likely require a change in the allocation of attentional resources in order to select a new path. Such an allocation of attention to select behaviourally relevant items amongst competitors, both externally (Worden, Foxe, & Wang, 2000) or from memory (Stokes, Atherton, Patai, & Nobre, 2012), has consistently been linked to a reduction in alpha band (8-12 Hz) power. Thus, we hypothesised that alpha power might reduce during Backtracking events compared to control events. Consistent with our prediction we indeed found a significant decrease in alpha power preceding the onset of Backtracking compared to Non) backtracking events (−700 to −150ms, p=.008), over a wide range of posterior sensors (Figure 3B). Control frequency bands (theta, 3-7 Hz; and beta, 15-25 Hz) did not reveal any effects in the pre-backtracking time-window. Moreover, the change in alpha activity could not be explained by differences in saccadic activity across the trial types (t(1,23)=−.9,p>.1). There was no correlation between the trial-wise alpha decrease and subsequent reaction times (i.e. time between button presses, as this was spontaneous behaviour) to choose to Back-track (r<0.03, p >.1). To confirm the robustness of our results, we also investigated the decreased alpha power in Backtracking in only those participants who had a minimum of five such events (n=21), and found the effect replicated in a similar time)window (−600 to −10ms, p=0.034). We also checked whether there was a modulation in alpha power before participants made any type of turn (i.e. left or right, but no backtracking), and found no significant effect, indicating that the process of backtracking selectively involves allocation of attentional resources and is not a confound of visual activity induced by direction changes. Additionally, when directly comparing backtracking to turning events, there was a significant decrease in alpha power in the pre-step time-window (p=0.005), further underscoring the notion that there are differences in attentional resource allocation during backtracking specifically.

In this study, we report three main findings: firstly, we show that visually identical changes in the environment are processed differently depending on whether they will aid reaching the goal (Shortcuts) or hinder it (False Shortcuts): Shortcuts induced changes in early event-related fields and showed increased theta power compared to False Shortcuts. Furthermore, False Shortcuts Towards the goal also showed increased theta power compared to False Shortcuts Away. As False Shortcuts Away can be distinguished on the basis of the goal direction whereas Shortcuts and False Shortcuts Towards required memory for the structure of the environment to distinguish them, the increased theta power at these latter events is consistent with the suggestions that theta may play a role in aiding future navigational planning through retrieval (Kaplan et al., 2014), imagery (Kaplan et al., 2017) and with the increased theta activity elicited when a longer distance was expected in the future than shorter one (Bush et al., 2017; Caplan et al., 2001; Vass et al., 2016). Alternatively the theta response may be more consistent with an increase in the conflict between choices or stimuli, as has been observed in previous navigation studies (Watrous, Fried, & Ekstrom, 2011; Weidemann, Mollison, & Kahana, 2009) and cognitive control paradigms (for review see Cavanagh & Frank, 2014). Finally, we explored the neural activity that occurred prior to spontaneous, internally generated back)tracking events, and observed a decrease in alpha power, consistent with changes in the allocation of attention (for review see Klimesch, 2012), which has been argued to be important during navigation (Grossberg & Pilly, 2013; Morris & Frey, 1997). Notably, a previous study also found decreased alpha power in relation to direction changes during passive virtual movement in navigation (Gramann et al., 2010). We show here that this suppression is also present for changes initiated by the person navigating and that these putative markers of attentional selection precede the onset of the direction change. Our results help extend our understanding of how the human brain responds to changes in the structure of environment during navigation and help provide much needed evidence to sculpt computational models of navigational guidance systems.

## Methods

### Participants

Twenty-five subjects (mean age: 22.5 ±3.9 years, range: 18-31; 12 female). Participants were administered a questionnaire regarding their navigation abilities/strategies (Santa Barbara Sense of Direction Scale; mean score=5.1, range: 3.2-6.8). All participants had normal to corrected vision, reported no medical implant containing metal, no history of neurological or psychiatric condition, color blindness, and did not suffer from claustrophobia. All participants gave written consent to participate to the study in accordance with the Birkbeck-UCL Centre for Neuroimaging ethics committee. Subjects were compensated with a minimum of £70 plus an additional £10 reward for good performance during the scan. One participant was excluded from the final sample because the MEG files were corrupted.

### VR environment: Lavaworld

A virtual island maze environment was created using Vizard virtual reality software (© WorldViz). The maze was a grid network, consisting of ‘sand’ areas that were walkable, and ‘lava’ areas, which were unpassable and as such were like walls in a traditional maze. However, the whole maze layout was flat, so there was visibility into the distance over both sand and lava. This allowed participants to stay oriented in the maze throughout the task. Orientation cues were provided by four unique large objects in the distance. Movement was controlled by 4 buttons: left, right, forwards and backwards. Pressing left, right or backwards moved the participant to the grid square to the left, right or behind respectively, and rotated the view accordingly. Similarly, pressing forward moved the participant to the next square along. See Figure 1 for a participant viewpoint at one point in the maze. Participants were tested over two days, on day 1 they were trained on the maze, and on day 2 they were tested in the MEG scanner.

#### Training

On the first day, participants were trained on the maze (25 × 15 grid) to find goal locations. During this phase, all goal objects (20 in total, distributed across the maze) were visible at all times, and participants navigated from one to the next based on the currently displayed target object (displayed in the top-right corner of the screen). After one hour of training, subjects were given a test to establish how well they had learnt the object locations. On a blank grid, where only the lava was marked, participants had to place all the objects they remembered. They were given feedback from the experimenter, and if needed, prompts as to the missing objects. This memory-test was repeated twice more during the training, after 1.5 and 2 hours. At completion, for participants to return for the MEG phase on the second day, they had to score at 100% accuracy in placing the objects.

#### Test & MEG scan

On the test day, participants were given a brief refresher of the maze with the objects. While in the MRI scanner, participants performed the test phase of the experiment. A single trial in the test phase is defined as being informed which is the new goal object, and then finding the way to, and arriving at, it. During the test phase, two things were different from training: 1) target objects were not visible, so participants had to navigate between them based on their memories of the locations, and 2) the lava could move, blocking some paths and creating new ones. During each journey to an object, one change occurred in the lava layout at a specific location. At the point of a change, the screen froze for 4 seconds to ensure that participants had an opportunity to detect the change and consider their path options. These changes could either be Detours (when a piece of lava was added to block the current path on the grid, thus forcing the participant to take an new, longer, route to their goal); Shortcuts (a piece of lava was removed and replaced with traversable sand, allowing the participant to pick a shorter route); False Shortcuts (visually identical to Shortcuts, but such that traversing them would increase the net distance to the goal because of the layout of the maze); and a Control condition in which the screen froze, but no lava was added/removed). False Shortcuts came in two classes: False Shortcuts Towards and False Shortcuts Away from the goal, depending on whether or not traversing them would appear to move closer to the goal. For Detours and Shortcuts, there were also two levels of change to the (optimal) new path, either 4 or 8 grid steps extra/less, respectively. See Figure 1 and S1 for example schematics of these changes. Finally, there were control ‘Follow’ trials, which started with an arrow that indicated the direction to travel. In this case, participants were required to follow the twists and turns of the arrow until a new target object appeared. The comparison of Navigation’ vs ‘Follow’ movements allowed us to relate our results to those of previous experiments (Howard et al., 2014; Javadi et al., 2017) (Howard et al., 2014; Javadi et al., 2017; Patai et al., 2017). Before scanning, participants were allowed to familiarize themselves with the scanner button pad, and with the changes that would occur. This involved presenting them with a novel environment that had not been experienced on day one, and which had no object, different distal cues and a different maze layout, to avoid any confounds or confusion with training and test mazes. Participants could then practice the task in this new environment, and accustom themselves to the controls (button pad with 4 active buttons: left, right, forward, and turn around) and to the appearance of changes to the lava.

### MEG Recording and Analysis

Recordings were made using a 275-channel Canadian Thin Films (CTF) MEG system with superconducting quantum interference device (SQUID)-based axial gradiometers (VSM Med)- Tech) and second)order gradients in a magnetically shielded room. Neuromagnetic signals were digitized continuously at a sampling rate of 480 Hz and then band-pass filtered in the 0.1–120 Hz range. Head positioning coils were attached to nasion, left, and right auricular sites to provide anatomical coregistration to a template brain. Preprocessing and analysis of MEG data was done using Fieldtrip (Oostenveld, Fries, Maris, & Schoffelen, 2011). Independent) component analysis (ICA) was performed on the continuous data, leading to the identification of blink, saccade and cardiac components, which were removed. MEG data were subsequently parsed into epochs starting 1000ms before, and ending 4000ms after, the onset of the change point. We also parsed the data around ‘backtracking’ events (-1000 to 1000ms), which were defined as steps on which participants turned around and thus spontaneously changed their paths. Specifically, a ‘Back-tracking’ event was defined by when participants pressed the backwards button and returned to a step along the route they had just come down. Non-back-tracking events were picked from the participants’ other paths and were matched to these in the relative number of steps taken before the back-tracking happened (for example: halfway through the route). We did not use the absolute number of steps, because trials that contained a back-tracking event were often much longer, and thus the step number at which a back-tracking event occurred could have already been at (or past) a goal point in a non-back-tracking trial. We used step matching rather than elapsed time matching because participants’ speed was not controlled and we wanted to match the events based on actual navigation steps and not potential time differences due to stationary behavior.

Given the exploratory nature of this study, we investigated effects of change type using all sensors, and we chunked the Detour and Shortcut epochs into 1000ms bins (resulting in 4 bins for the whole 4 second change period). Here we report significant effects found, cluster corrected for multiple comparisons. For time)frequency analyses, we used the same exploratory method, but specifiying the frequency ranges based on *apriori* bands as previously reported in the literature. We also combined both lengths of Shortcut (−4 / −8) for comparison with False Shortcuts.

### Experimental Design and Statistical Analysis

We used a 2 × 2 within subjects design, with two levels of change type (Detour or Shortcut) and two levels of magnitude (long: 8 steps, or short: 4 steps). In addition, we included False Shortcuts to ensure participants had to be selective about which new openings in the paths they would choose. Participants performed a total of 120 routes, with one change occurring in each route. Each route started from a previous goal and ended at the new goal object for that trial. We used repeated-measures ANOVAs to test for behavioural differences (accuracy (shortest path chosen or not), extra steps, back-tracking trials) between conditions. We also calculated d-prime and criterion (signal detection theory measures) to quantify the bias to take a False Shortcuts Towards instead of Away from a goal (both false alarms calculated relative to correct shortcuts, which are hits). We recorded the response time to make the first choice after the 4 seconds elapsed, but due to the 4 second delay, we do not interpret this as a traditional decision-making reaction time. We excluded Control (i.e Freeze) events from all subsequent analyses, as it was a control condition and participants had very low accuracy (74% correct). Post-test debriefing indicated that this was likely due to participants finding it confusing that despite the screen freezing for 4 sec there was no apparent change and thus they changed their route choice in case they had missed a change. Additionally, given the limit on trial numbers, we were not able to investigate differences between correct rejections of False Shortcuts and those that were taken mistakenly.

A ‘Back-tracking’ event was defined by when participants pressed the backwards button and returned to a step along the route they had just come down. Non-back-tracking events were picked from the participant’s other paths and were matched to these in the relative number of steps taken before the back-tracking happened (for example: halfway through the route). We did not use the absolute number of steps, because trials that contained a back-tracking event were often much longer, and thus the step number at which a back-tracking event occurred could have already been at (or past) a goal point in a non-back-tracking trial. We used step matching rather than elapsed time matching because participants’ speed was not controlled and we wanted to match the events based on actual navigation steps and not potential time differences due to stationary behavior.

To analyse the MEG data we focused on event-related fields (ERFs), as well as time-frequency analysis. However, due to the nature of the task (free-viewing during navigation), and despite the ICA correction, we were unable to exclude fully the possibility that some oscillatory signatures would be contaminated by eye-movements. We therefore looked at the difference between the saccade variance as measured by ICA across different conditions, and found that the only comparison that was not potentially confounded by eye-movement was the backtracking vs non-backtracking comparison (see Supp. Results for details).

## Acknowledgements

We thank Gareth Barnes, UCL WCHN, for helpful comments on the analysis and Mate Lengyel, University of Cambridge, for advice on the experimental design. This work was supported by the Wellcome Trust (grant 094850/Z/10/Z) and James S. McDonnell Foundation to H.J.S, and the Gatsby Charitable Foundation (P.D.). The authors declare no competing financial interests. P.D. is currently on sabbatical at Uber AI Lab.

## Supplemental Results

For completeness, we include discussion of other analyses of the changes in the environment. However, due to eye-movements, we cannot explicitly exclude the possibility that effects are contaminated by differences in saccadic behaviour between conditions (as measured by variance in the saccade components derived from the ICA).

### Event-Related Fields

#### Detours vs Shortcuts

Two significant time periods emerged from our analysis, 400-600ms and 700-1000ms after the onset of the terrain change. In both of these, there was a larger deflection for Shortcuts than Detours: an earlier left fronto-temporal effect, followed by a right, temporal)occipital effect (see Figure S1B). However, the saccade variance was significantly different between conditions (paired)t-test, t(1,23)=2.5,p=0.02).

#### Large vs Small Changes

To investigate neural responses related to the magnitude of the change in path instigated by the terrain change point, we combined long Detours and Shortcuts (+/−8) and short Detours and Shortcuts (+/−4). We found a significant frontal effect from 250-800ms, with changes that induced a large change in path showing a larger deflection. However, upon inspection of the topography and the ICA variance (paired-t-test, t(1,23)=4.9,p<0.001), we believe that this effect could be explained by eye-movement contamination.

#### Interaction between Change Type and Magnitude

A repeated measures ANOVA revealed no significant effect of change type, magnitude and no interaction. There was however a significant effect of eye) movements between conditions (type, F(1,23)=6.4,p<0.019, magnitude F(1,23)=24.02,p<0.001, interaction F(1,23)=64.04,p<0.001), with Detours (+8) showing the largest variance compared to all conditions (all t(1,23)>4.1, p<0.001).

**Table S1:**
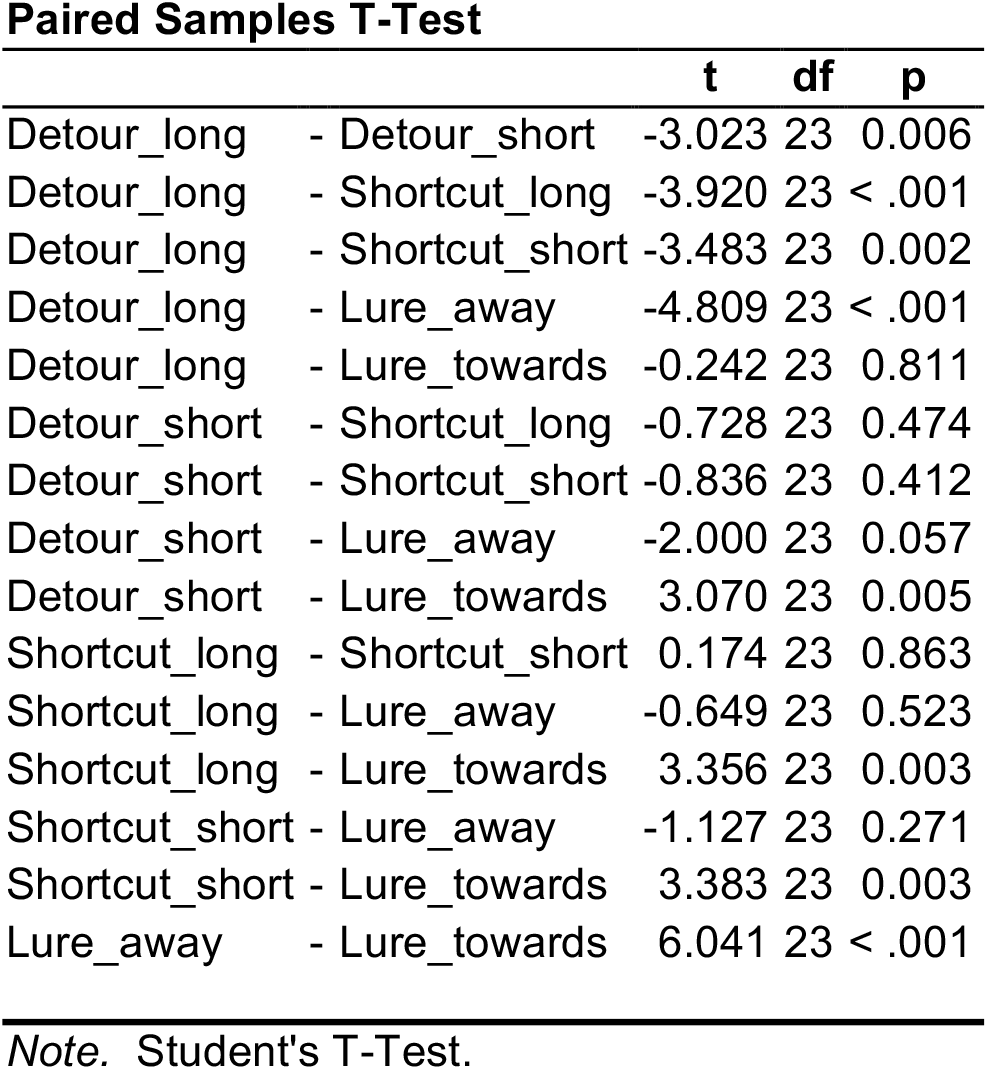
Accuracy

**Table S2:**
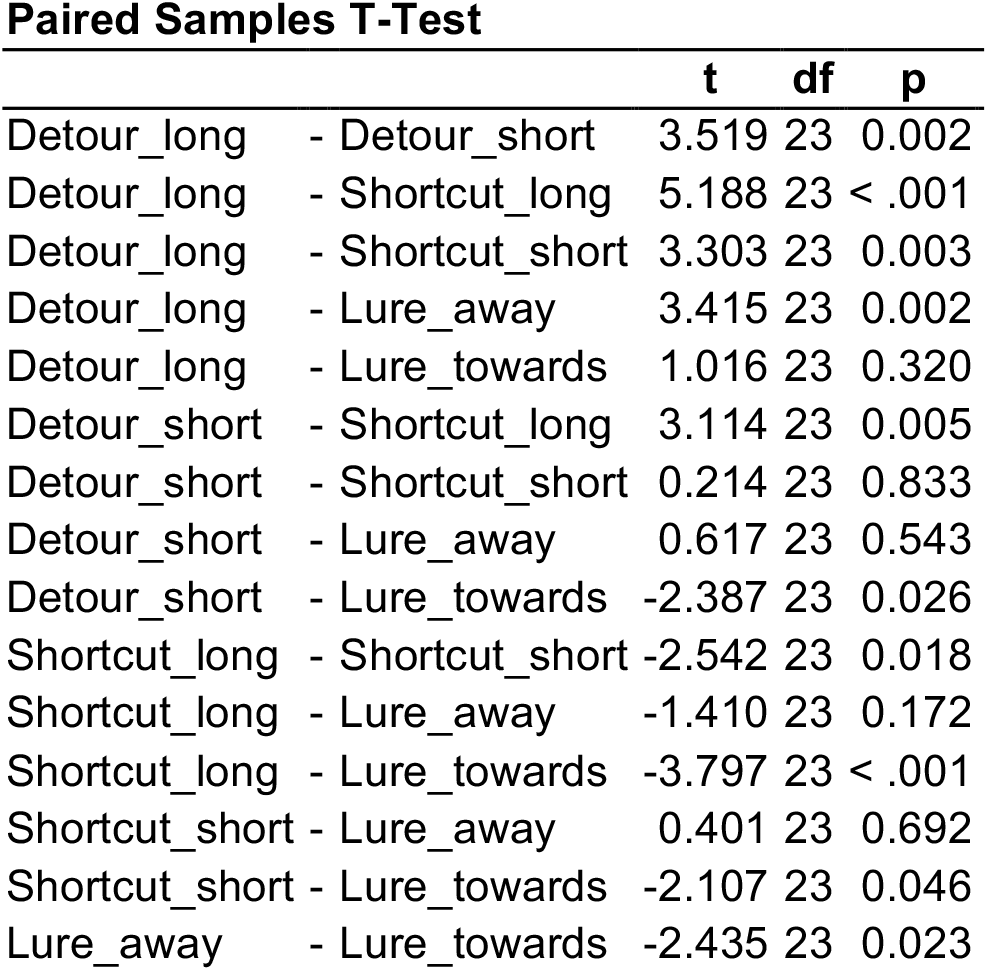
Backtracking

**Figure S1.**
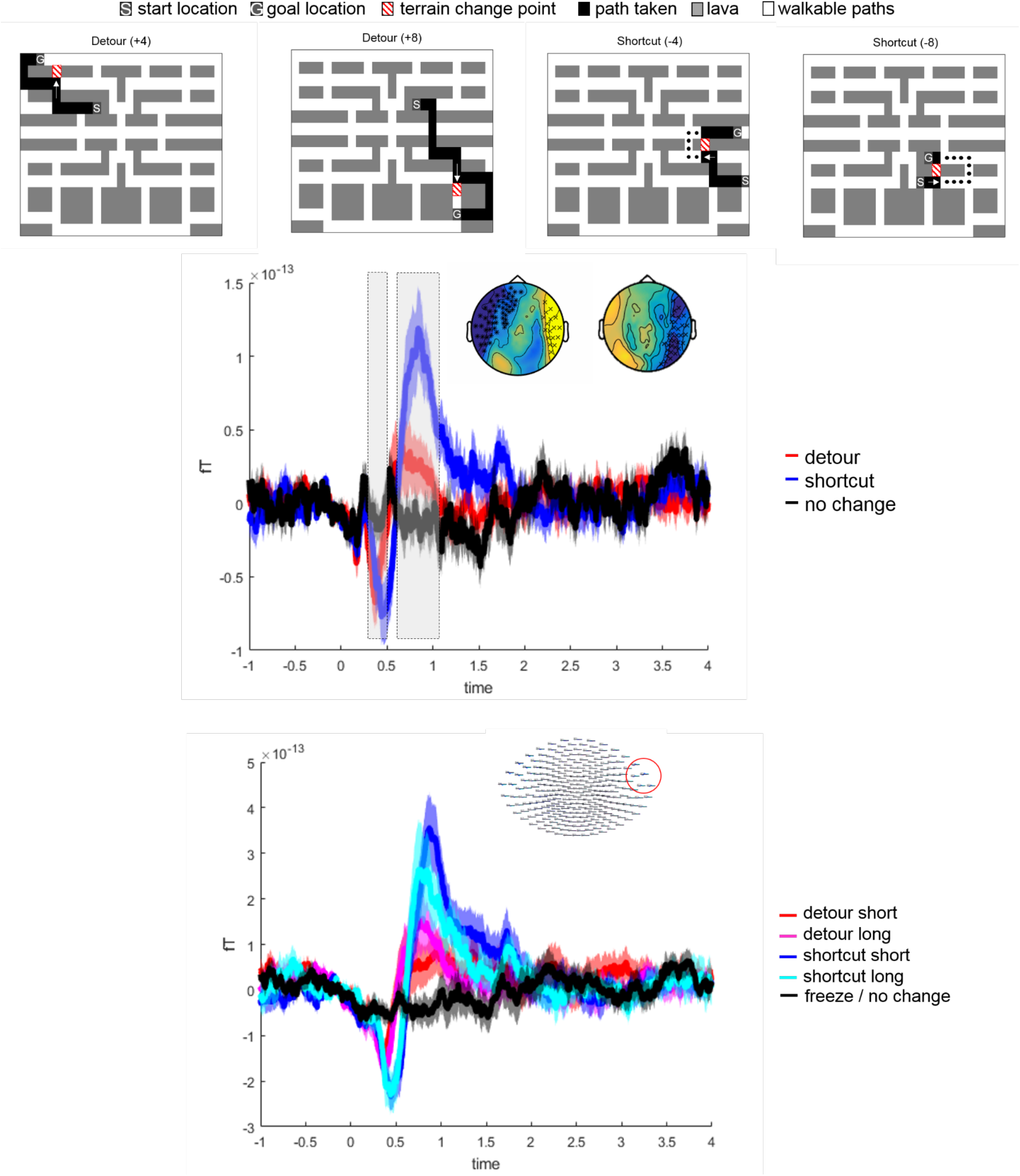
Even-related fields to Detours and Changes in Magnitude of Novel Path. A) Example of different change types, including Detours. The arrow marks the path the Participant would have taken if the path had not been blocked. B) Detours vs Shortcuts Showed significant hanges in evoked fields around 400 ms after change onset (Over magnitude of change). C) Effect plotted for sensors shown in top right corner, for all Detour and Shortcut conditions showing that the size of the change (=/− 4 vs 8 steps).

